# Quality Control and Integration of Genotypes from Two Calling Pipelines for Whole Genome Sequence Data in the Alzheimer’s Disease Sequencing Project

**DOI:** 10.1101/318857

**Authors:** Adam C. Naj, Honghuang Lin, Badri N. Vardarajan, Simon White, Daniel Lancour, Yiyi Ma, Michael Schmidt, Fangui Sun, Mariusz Butkiewicz, William S. Bush, Brian W. Kunkle, John Malamon, Najaf Amin, Seung Hoan Choi, Kara L. Hamilton-Nelson, Sven J. van der Lee, Namrata Gupta, Daniel C. Koboldt, Mohamad Saad, Bowen Wang, Alejandro Q. Nato, Harkirat K. Sohi, Amanda Kuzma, Alzheimer’s Disease Sequencing Project (ADSP), Li-San Wang, L. Adrienne Cupples, Cornelia van Duijn, Sudha Seshadri, Gerard D. Schellenberg, Eric Boerwinkle, Joshua C. Bis, Josée Dupuis, William J Salerno, Ellen M. Wijsman, Eden R. Martin, Anita L. DeStefano

**Affiliations:** Department of Biostatistics and Epidemiology, and; Department of Pathology and Laboratory Medicine, Perelman School of Medicine, University of Pennsylvania, Philadelphia, PA, USA; Department of Medicine, Boston University School of Medicine, Boston, MA, USA; Department of Neurology, Columbia University Medical Center, New York, NY, USA; Human Genome Sequencing Center, Baylor College of Medicine, Houston, TX, USA; Department of Biomedical Genetics, Boston University School of Medicine, Boston, MA, USA; John P. Hussman Institute for Human Genetics, University of Miami Miller School of Medicine, Miami, FL, USA; Department of Biostatistics, Boston University School of Public Health, Boston, MA, USA; Department of Epidemiology and Biostatistics, Case Western Reserve University, Cleveland, OH, USA; Department of Epidemiology, Erasmus Medical Center, Rotterdam, The Netherlands; Medical and Population Genetics Program, Broad Institute, Cambridge, MA, USA; Institute for Genomic Medicine, Nationwide Children's Hospital, Columbus, OH, USA; Department of Biostatistics, and; Division of Medical Genetics, and; Department of Statistics, University of Washington, Seattle, WA, USA; The Framingham Heart Study, Framingham, MA, USA; Department of Neurology, Boston University School of Medicine, Boston, MA, USA; Human Genetics Center University of Texas Health Science Center, Baylor College of Medicine, Houston, TX, USA; Cardiovascular Health Research Unit, Department of Medicine, University of Washington, Seattle, WA, USA

**Keywords:** quality control, whole genome sequencing, Atlas, GATK, Mendelian inconsistencies, consensus calling

## Abstract

The Alzheimer’s Disease Sequencing Project (ADSP) performed whole genome sequencing (WGS) of 584 subjects from 111 multiplex families at three sequencing centers. Genotype calling of single nucleotide variants (SNVs) and insertion-deletion variants (indels) was performed centrally using *GATK-HaplotypeCaller* and *Atlas V2*. The ADSP Quality Control (QC) Working Group applied QC protocols to project-level variant call format files (VCFs) from each pipeline, and developed and implemented a novel protocol, termed “consensus calling,” to combine genotype calls from both pipelines into a single high-quality set. QC was applied to autosomal bi-allelic SNVs and indels, and included pipeline-recommended QC filters, variant-level QC, and sample-level QC. Low-quality variants or genotypes were excluded, and sample outliers were noted. Quality was assessed by examining Mendelian inconsistencies (MIs) among 67 parent-offspring pairs, and MIs were used to establish additional genotype-specific filters for *GATK* calls. After QC, 578 subjects remained. Pipeline-specific QC excluded ~12.0% of *GATK* and 14.5% of *Atlas* SNVs. Between pipelines, ~91% of SNV genotypes across all QCed variants were concordant; 4.23% and 4.56% of genotypes were exclusive to *Atlas* or *GATK*, respectively; the remaining ~0.01% of discordant genotypes were excluded. For indels, variant-level QC excluded ~36.8% of *GATK* and 35.3% of *Atlas* indels. Between pipelines, ~55.6% of indel genotypes were concordant; while 10.3% and 28.3% were exclusive to *Atlas* or *GATK*, respectively; and ~0.29% of discordant genotypes were. The final WGS consensus dataset contains 27,896,774 SNVs and 3,133,926 indels and is publicly available.

**Abbreviations:** *AD*, Alzheimer’s disease; *QC*, Quality Control; *LSSAC*, Large-Scale Sequencing and Analysis Center; *Broad*, Broad Institute Genomics Service; *Baylor*, Baylor College of Medicine Human Genome Sequencing Center; *WashU*, Washington University-St. Louis McDonnell Genome Institute; *WGS*, whole genome sequencing; *WES*, whole exome sequencing; *indel*, insertion-deletion variants; *VCF*, variant control format; *MI*, Mendelian inconsistency; *MC*, Mendelian consistency; *GWAS*, genome-wide association study; *VR*, referent allele read depth; *DP*, overall read depth; *MS*, mapping score; *GQ*, genotype quality score; *Ti/Tv*, Transition/Transversion; *CS*, concordance code

## Introduction

While genome-wide association studies (GWAS) have successfully identified thousands of common genetic variants associated with hundreds of complex diseases and traits, next-generation sequencing (NGS) technologies aim to widen the scope and ability of genomic studies to identify the genetic underpinnings of disease. Unlike array-based GWAS genotyping, NGS technologies, which include whole genome sequencing (WGS) and whole exome sequencing (WES), are able to comprehensively capture genotypes for both common and rare single nucleotide variants, insertion-deletion polymorphisms, and even structural variants that may contribute to disease risk. By capturing all sequence within targeted regions, NGS may facilitate identification of causal variants rather than merely associated variants. It achieves a substantial gain in base-pair coverage of the genome compared to high-density GWAS array genotyping, a multi-fold increase in read depth compared to traditional Sanger sequencing, and a much lower per-base cost.^1^

Both WGS and WES genotype calling can be affected by a variety of types of errors or sources of bias, such as sample swaps and low call rates that can alter the quality of genotypes used in analyses and thus affect the ability of analyses to detect associations with disease.

Quality issues that are unique to sequencing assays can arise at multiple steps in the process. During the library preparation and sequencing phases, these include low quality reads resulting from duplications, unfavorable base composition of the amplified sequence, inclusion of tag (adapter/barcode) sequences into reads; as well as read contamination from external sources, such as bacteria in DNA samples used for sequencing.^2^ At the alignment phase, quality issues can arise from alignment to duplicated genomic regions or repeat-rich regions, and require assessments of read depth, mapping quality, insert size and the number of discordantly mapped paired reads^3^. Issues at these phases can only be remedied by appropriate experimental design and use of sensitive bioinformatic tools. At the variant-calling level, quality issues include elevated numbers of novel non-synonymous SNPs or excesses of ‘private’ variants (within individual samples), allelic read ratio biases (causing true homozygotes to be called as heterozygotes), and low-quality calls based on low read depths. While a number of QC software packages and protocols exist for the cleaning at the raw data phase^2,4,5^ and at the alignment phase,^6^ few tools or protocols exist^7^ at the variant-calling/post-variant-calling phase. For this reason, a novel QC protocol, including the development of a consensus-calling approach to integrate genotype calls from multiple pipelines, was developed for WGS data in the Alzheimer Disease Sequencing Project (ADSP).

The ADSP is a collaboration between the National Institutes on Aging (NIA) and the National Human Genome Research Institute (NHGRI), with data contributions from the Alzheimer Disease Genetics Consortium (ADGC) and the neurology working group of the Cohorts for Heart and Aging Research in Genomic Epidemiology (CHARGE) consortium, and sequencing contributions from the Large Scale Sequencing and Analysis Centers (LSSACs) at Baylor University, the Broad Institute and Washington University-St. Louis. The ADSP was initiated to identify both protective and risk genetic variants for Alzheimer Disease (AD) [MIM: 104300], a devastating neurodegenerative disorder characterized by progressive loss of cognitive function. The ADSP generated WGS on 584 individuals from 111 large, multiplex late onset AD families to detect risk variants having large effect on familial forms of AD. Raw data processing, map alignment, and variant calling were performed by both the Broad Institute, which called variants using the GATK-HaplotypeCaller package,^8-10^ and the Baylor College of Medicine Human Genome Sequencing Center, which called variants using the Atlas V2 pipeline.^11^

The ADSP QC protocol integrated QC strategies from multiple sources including the CHARGE consortium QC protocol,^12^ prior sequencing study experiences of ADGC and CHARGE investigators, GWAS QC approaches, and proprietary QC recommendations for the Atlas V2 and GATK-HaplotypeCaller genotype calling pipelines. The final protocol included 1) independent QC of each variant calling set, 2) comparison of the QCed calls from each pipeline, and 3) implementation of a consensus calling protocol to remove discordant calls and integrate genotypes called in only one pipeline, resulting in a single set of SNV and indel genotypes for analysis.

Here we discuss the workflow and implementation of this novel QC protocol and consensus calling approach on WGS autosomal SNV and indel data in the ADSP. We demonstrate improved quality of SNV and indel genotype data after pipeline-specific QC filtering and show further improvements in quality through higher concordance of genotype calls and lower rate of Mendelian inconsistencies by combining genotype calls from two calling pipelines compared to using calls from only one pipeline.

## Subjects and Methods

### Subjects

The ADSP family study spans seven cohorts. Detailed description of the study design has been published elsewhere.^13-15^ In brief, 1,100 multiplex AD families were screened to identify 111 high priority families consisting of more than three AD cases with limited presence of the *APOE*ε4 allele and other known pathogenic variants (e.g., *APP, PSEN1/2)*. All subjects selected for sequencing had either genome-wide or exome chip SNV genotype data available.

A total of 584 subjects were selected for sequencing from these pedigrees. Three subjects were sequenced in replicate at all three sequencing centers, adding six more samples, for a total of 590 samples. Six samples were dropped because of sequencing quality issues, including one with low DNA concentration, three with poor GWAS concordance, and two with low-quality sequence data. Of the remaining 584 samples from 578 unique subjects, 12 samples were resequenced due to issues with reagents. These family data included individuals of European American, African American, and Caribbean Hispanic ancestry, and members of a large multi-generational pedigree with high burden of AD from a Dutch isolate^16^ (Table S1).

### Whole genome sequencing methods

Genomic DNA from whole blood, frozen brain, or fibroblasts was sent to one of three LSSACs: Broad Institute Genomics Service (Broad), Baylor College of Medicine Human Genome Sequencing Center (Baylor), and McDonnell Genome Institute at the Washington University in St. Louis (WashU). The breakdown of samples sequenced by each of three centers is shown in Table S1. Illumina WGS technology was used at all three centers. Library preparation and sequencing protocols details are provided in the Supplementary Materials (Text S1; Table S2).

### Whole genome alignment and variant genotype calling

After sequencing, Broad and Baylor performed alignment and variant calling on all whole genomes from all three LSSACs. Genome alignment at Broad was performed using the 1000 Genomes version of the GRCh37/hg19 build, while alignment at Baylor was performed using the GRCh37-lite version. Broad and Baylor subsequently applied two variant genotype callers, GATK-HaplotypeCaller V2.6 and Atlas V2, respectively, to the sequencing data. Additional details regarding genome alignment are provided in Table S2, while a detailed description and workflow characterizing variant genotype calling in GATK and Atlas are described in Text S2.

### Pre-QC sample checking

Concordance checking between high-density genotyping chip-based genotype data (either GWAS or exome chip) was performed by the sequencing centers prior to implementing the QC protocol. Samples with a concordance rate of <80% between chip genotype data and sequence were removed from subsequent QC and analysis. The final QCed genotype set includes only a single set of sequence genotypes for the three replicated subjects, specifically the data generated by the sequencing center to which other members of the subjects’ pedigrees were originally allocated. Finally, only data for subjects that were successfully called in both Atlas and GATK pipelines were carried forward to QC.

Familial relationships between samples were evaluated via the software suite PBAP (Pedigree-based analysis pipeline)^17^ using selected sets of ~5,000-6,000 high frequency (MAF>0.05) variants from the GWAS arrays to obtain maximum likelihood estimates of pairwise IBD sharing coefficients. Relative pairs with coefficient estimates that deviated strongly from expectation based on the reported familial relationships were flagged but not excluded prior to QC.

Checking for Mendelian inconsistencies (MIs) between related individuals was performed in 67 parent-offspring pairs that were sequenced within the ADSP pedigrees using Pedcheck software or R scripts.^18^ This was done prior to QC to establish baseline rates of MIs within each pipeline and after QC in order to quantify the improvement in data quality following the implementation of QC filters. Given that only biallelic variants were considered and no trios (with both parents) were available, MIs could only be detected when the parent and offspring were homozygotes for different alleles (depicted in Figure S1). Because the incorrect genotype in each pair could not be established, genotypes involved in MIs were not excluded from QCed datasets.

### Pipeline-specific quality control

SNV and indel genotypes were QCed independently, although the same protocol was applied to each variant type. QC was applied to biallelic autosomal SNVs and indels; any variants with more than two alleles were removed from the data set prior to other QC steps and are not addressed in the current study. Three levels of QC were applied: (a) pipeline-specific ‘primary’ QC (i.e., sequencing center-recommended filtering); (b) standardized variant-level QC; and (c) sample-level QC.

Figure 1 depicts all components of the QC protocol. For pipeline-specific primary QC, Atlas V2 genotype calls were first evaluated at a genotype-level. Genotypes that had a low read depth (DP <10) or an out-of-range allelic read ratio (VR/DP ≤0.75 or VR/DP ≥0.25 for heterozygous genotypes, where VR and DP are referent allele and overall read depths, respectively) were set to missing. Next on the variant level, any variants with low mapping score (MS) (in the VCF’s “INFO” field, MS <0.8) and any variants with completely missing genotypes were flagged/excluded as failed variants. GATK-HaplotypeCaller primary QC was applied at the variant level, excluding any variants that were not flagged as “PASS” by the VQSR algorithm, thus excluding variants outside the 95% sensitivity tranches (i.e., the lowest 5% of recalibrated quality scores).

**Figure 1.**
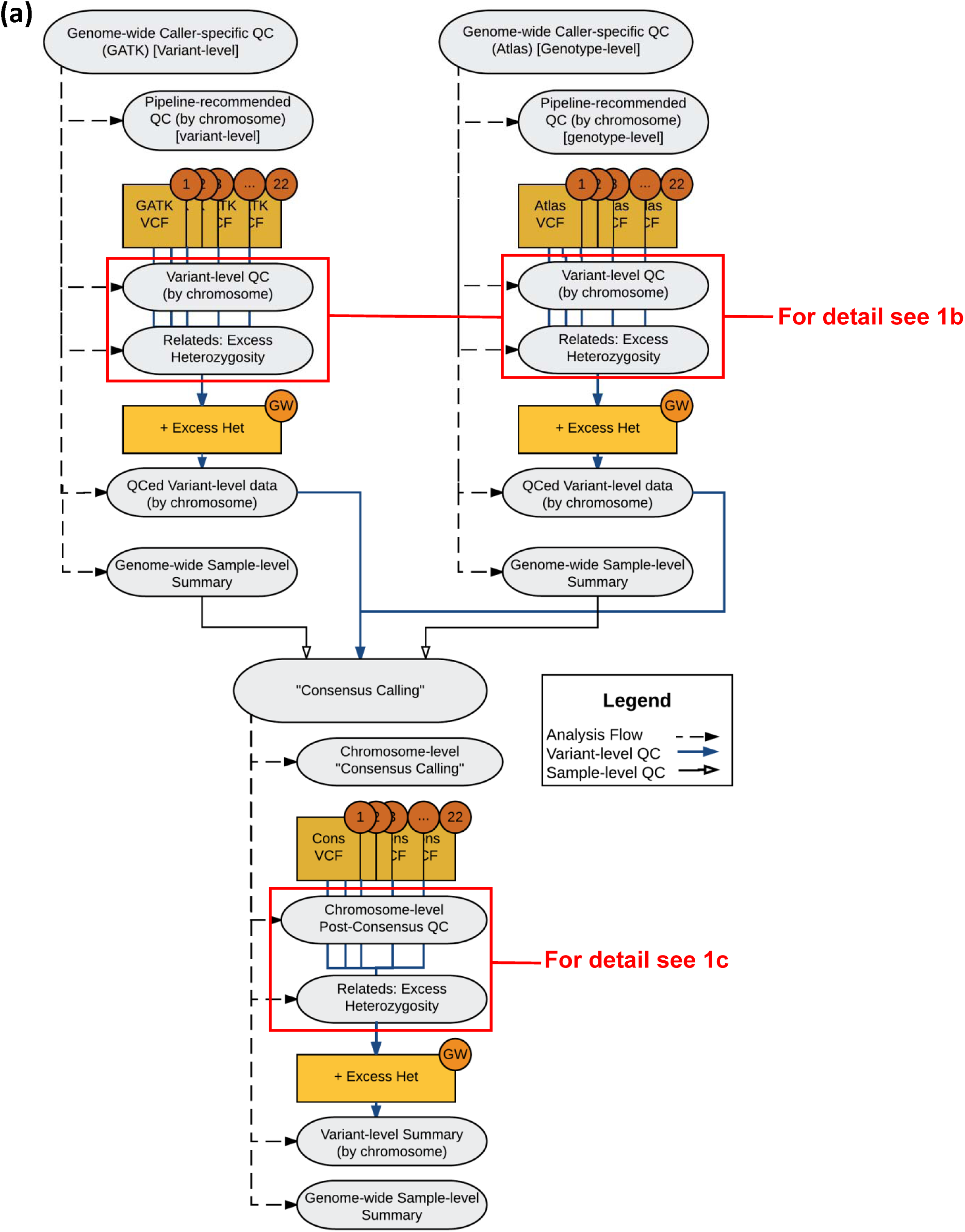
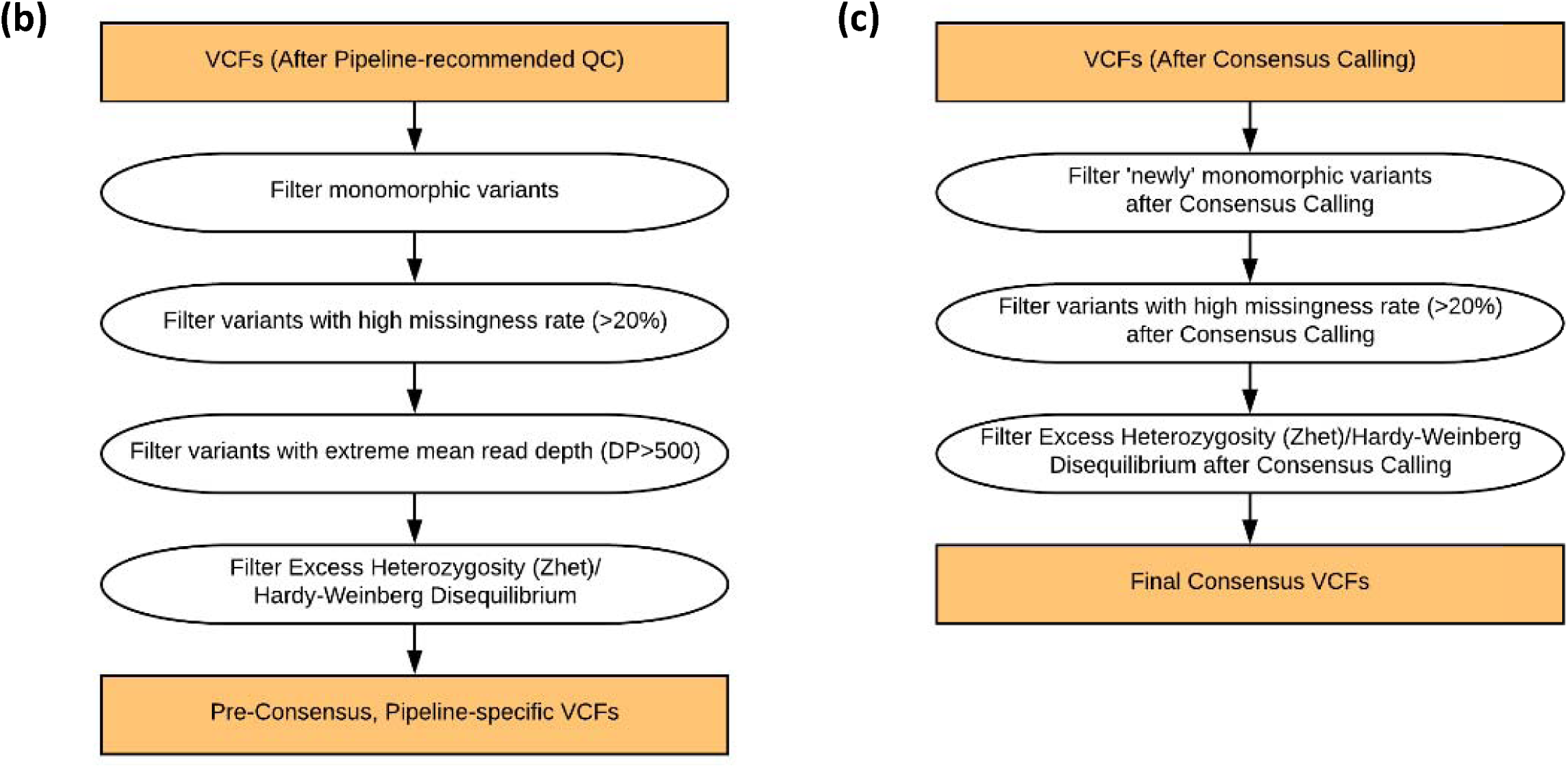
Diagrams of the Caller-specific QC and Consensus Calling Pipeline, including **(a)** an overview diagram of the process, **(b)** details of the caller-specific variant-level QC steps, and **(c)** details of the post-consensus variant-level QC steps.

After primary QC, variant-level QC was applied to the remaining variants in the Atlas and GATK VCFs. Variant filters (exclusions) were applied in the following order: 1) monomorphic; 2) high missing rate (≥20%); 3) high read depth (>500 reads); and 4) extreme heterozygosity (>5 SD from mean z-score across all variants). For step #4, we evaluated excess heterozygosity empirically in lieu of testing for departure from Hardy-Weinberg equilibrium (HWE) due to concerns about the potentially biasing effects of familial relationships on variance estimation and consequently on the *P*-value. Instead, a score statistic approach was applied across all sequenced individuals except members of the Dutch Isolate pedigrees, and this approach is described in the Supplement (Text S3).

After implementation of variant-level QC, within-individual quality metrics were estimated, and the distributions of these metrics were assessed to exclude potential outliers. Among these metrics were: (1) counts of singleton/doubleton variant calls to identify an excess of private variants; (2) genotype missingness rate (per individual) > 0.2; (3) Transition/Transverion (Ti/Tv) ratio outliers [for SNVs only]; (4) heterozygosity-to-homozygosity ratio (across all within-individual genotypes); and (5) mean read depth (across all within-individual genotypes). Samples with genotype missingness rate >0.2 were excluded and samples were examined as potential outliers if their values for any of the other criteria were greater than 6 SD from the mean value based on ethnic group. Three groups were defined: European American, Dutch Isolate, and Caribbean Hispanic

### Concordance matching

Once primary and variant-level filtering were completed, genotypes were compared between the two QCed VCFs. Table 1 contains the concordance codes (“CS”) derived for comparing the genotypes. In addition, a concordant set of genotype calls in which only genotypes present and concordant between the two QCed calling pipelines was created.

**Table 1.** Genotype concordance counts after preliminary and pipeline-specific QC for all possible genotype pairs between Atlas and GATK, across all individuals and variants. Included is a two-digit “concordance code” label, summarizing genotype pairs between the pipelines, Atlas (the first digit) and GATK (the second digit). Digit values are “0” for referent allele homozygote, “1” for heterozygote”, “2” for alternate allele homozygote, and “9” for missing/not available. The category “33” represents all genotypes (including referent homozygotes concordant between the two VCFs) for variants in which a different alternative allele was called between the two sets of VCFs.

| Code | Interpretation | Whole Genome Sequence (WGS) |  |
| --- | --- | --- | --- |
|  |  | SNV genotypes | Indel genotypes |
| 00 | Concordant reference allele (REF) homozygote | 12,822,511,444 | 1,020,985,877 |
| 11 | Concordant heterozygote | 1,193,810,170 | 75,163,355 |
| 22 | Concordant alternate allele (ALT) homozygote | 684,475,930 | 44,941,526 |
| 01 | Discordant; Atlas REF homozygote; GATK heterozygote | 247,853 | 1,662,092 |
| 02 | Discordant; Atlas REF homozygote; GATK ALT homozygote | 11,257 | 265,112 |
| 09 | Atlas REF homozygote; GATK missing | 647,757,411 | 200,377,988 |
| 10 | Discordant; Atlas heterozygote; GATK REF homozygote | 629,047 | 1,344,533 |
| 12 | Discordant; Atlas heterozygote; GATK ALT heterozygote | 103,484 | 2,317,666 |
| 19 | Atlas heterozygote; GATK missing | 24,279,157 | 7,477,521 |
| 20 | Discordant; Atlas ALT homozygote; GATK REF homozygote | 21,765 | 13,036 |
| 21 | Discordant; Atlas ALT homozygote; GATK heterozygote | 716,059 | 260,906 |
| 29 | Atlas ALT homozygote; GATK missing | 11,916,104 | 4,124,707 |
| 90 | Atlas missing; GATK REF homozygote | 619,012,699 | 516,306,709 |
| 91 | Atlas missing; GATK heterozygote | 87,373,375 | 43,251,176 |
| 92 | Atlas missing; GATK ALT homozygote | 31,581,133 | 14,675,561 |
| 99 | Atlas missing; GATK missing | 42,738,514 | 104,183,397 |
| 33 | All genotypes for variants with different alternative alleles between Atlas and GATK | 117,912 | 14,169,670 |

### Consensus Calling

The ‘concordant’ dataset contains high-certainty genotype calls that were identical in the two variant calling pipelines, but it did not allow inclusion of high-quality variants called in only one pipeline. To arrive at a single set of genotype calls that includes high-quality variants from each of the two variant calling pipelines, several approaches for consensus calling were explored. While methods for QC and reconciliation of variant calls from multiple calling algorithms have been developed and evaluated elsewhere,^19-21^ these methods did not enable the inclusion of certain filtering criteria we sought to implement and did not generate QC annotation at the variant-level that preserved all metrics from multiple callers. Additionally, they were not yet extensible implemented for indel variants, and to adapt these pipelines to perform these functions would have required extensive modifications or substantial additional scripting. The ADSP protocol described here was implemented on two platforms: first via R and Perl scripting in Linux for all SNV data, and then via SQL commands within a Hadoop Hive database system for indels. The steps of the consensus calling protocol are detailed in Box 1.

#### Box 1.

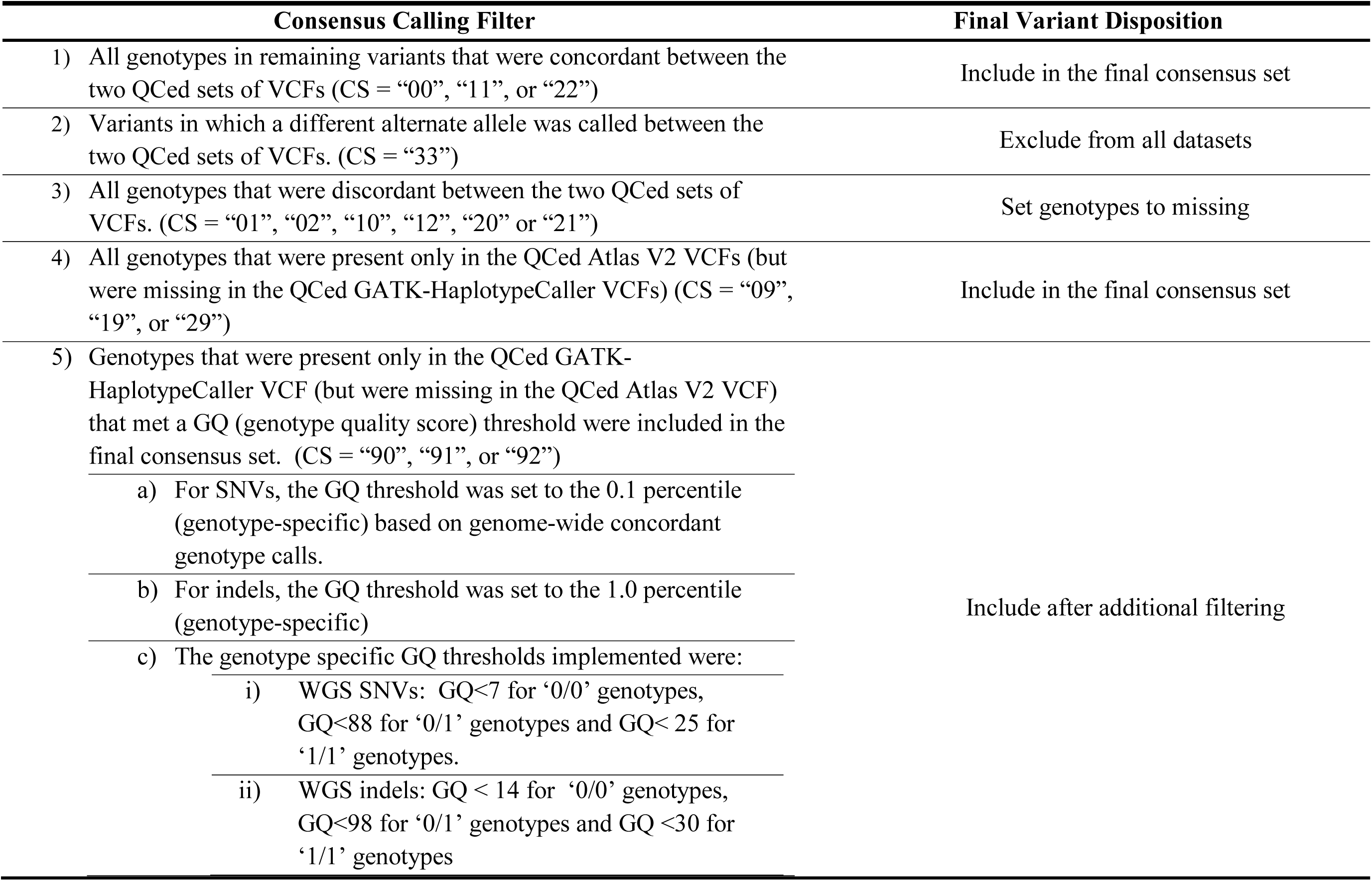
Steps and implementation hierarchy of the consensus calling process. Concordance code (“CS”) definitions are detailed in Table 1.

Examination of MIs in the genotype data after pipeline-specific QC identified a pattern of higher MI rates among variants called only in GATK compared to those called only in Atlas and those called in both pipelines, suggesting that additional genotype-level filtering for these variants was warranted. As exploration of genotype-specific QC metrics in the ADSP data identified GQ (“Genotype Quality”) as an informative filter to reduce MI rate, GQ filtering thresholds were implemented to remedy the higher rate of MIs in GATK data. The use of GQ as a filter here is supported by several prior studies that identified the utility of GQ in reducing MI rate^22,23^ and improving the quality of other metrics.^24-26^ Analyses to identify the genotype-specific GQ exclusion thresholds characterized are described in the Supplement (Text S4).

### Post-consensus calling variant-level QC

After the consensus calling process, variant-level QC filters were applied again to remove variants that now failed these criteria after consensus calling. These included the previous variant-level QC filtration steps: 1) excluding monomorphic variants; 2) excluding variants with missing rate ≥20%; and 3) excluding variants with extreme heterozygosity. After this final variant-level QC, individual sample statistics were also recomputed to assess quality after consensus calling.

### Post-processing

QCed data were reformatted as both PLINK binary-format files (*.bed, *.bim, *.fam) and as annotated VCFs, all containing only QC-passing variants. VCF releases included QC annotation (Table S3) on all variants indicating their final disposition (whether they were dropped in pipeline-specific QC, failed consensus calling or post-consensus QC, or whether they passed all QC stages).

### Evaluation of post-QC data quality through full pedigree Mendelian inconsistency evaluation

A summary measure from family-based genotype imputation was used to evaluate the variant-call pipelines. The full pedigree structures coupled with the existing GWAS data allowed use of imputation success of the WGS data in the context of GWAS markers as a proxy for genotype-call quality. This extended our ability to check for genotype quality beyond the limited case of parent-offspring pairs, which can identify errors only when there is homozygosity for different alleles in the two individuals (the parent-offspring pairs). The concept underlying this approach to genotype imputation is that, with the exception of rare de-novo mutations, real variants are inherited, and therefore high-quality called genotypes should show greater Mendelian consistency (MC) with pedigree inheritance vectors (IVs)^27^ than lower-quality called genotypes. We computed the MC probabilities within pedigrees and averaged these across pedigrees to obtain the “Imputation Rate” for each WGS position. Higher values of the imputation rate are indicative of higher quality called genotypes. The imputation rate was estimated by sampling IVs at the positions of each of the variants, conditional on the complete pedigree structure and sampled IVs at the positions of SNPs from the GWAS panels, followed by pedigree-based imputation^28^ from the WGS data. Details of the computation are provided in the Supplement (Text S5/Table S4). We computed the imputation rate for each of four categories of variants: (1) the 25,531,054 variants that initially passed QC in both pipelines, keeping the “concordant-only” genotypes at these variant positions [all non-concordant genotypes set to missing]; (2) the “consensus-only” 2,365,720 variants that passed QC in either the Atlas or GATK pipelines but not both; (3) the 1,241,253 variants that passed QC in only the Atlas pipeline; and (4) the 1,124,467 variants that passed QC in only the GATK pipeline. These last two categories represent each of the mutually exclusive variant sets that together represent the “consensus-only” category. We focused our evaluation on averages across all pedigrees with exactly *h* observed heterozygotes per pedigree, with h=1-4. This controls for variant allele frequency, which affects ease of successful family-based imputation because imputation is easier when there are fewer heterozygotes in a pedigree.

### ADSP QC Pipeline Availability

The ADSP QC Pipeline is implemented in Perl and runs on the command line, with parameters of the QC run, including filtering thresholds, input/output directories, and reporting rules specified in a separate “control” (parameter) file. The Perl script, sample control file, sample ‘fam’ file, and accompanying documentation, are all available at https://www.niagads.org/adsp/ADSP_QC Script. All formatting modifications performed on the VCF files during the QC process have been documented in the updated VCF header, ensuring the readability of these files by any program using the VCF v4.0 format.

## Results

### Comparisons of replicate samples across sequencing centers

The average pair-wise concordance rate between replicate samples prior to any QC, for the three replicates sequenced at all three centers, was 99.49% (ranging from 99.46% to 99.53%) for the GATK pipeline and 98.60% (ranging from 98.54% to 98.71%) for the Atlas pipeline. All three subjects showed similar patterns of between-replicate concordance, suggesting that the discrepancy of average concordance rates might be attributable to differences in calling pipeline rather than DNA quality. For comparison, we also examined the genotype concordance rate between two different unrelated subjects and found the average concordance rate was 91.52% for the GATK pipeline and 90.62% for the ATLAS pipeline. Both were significantly lower than the concordance rate of the same subject as expected. All comparisons of replicates are described in Text S6, and concordance counts are shown in Tables S5 and S6.

### WGS SNV and Indel QC on 578 family-based samples

The vast majority of QCed genotype calls, ~91%, were concordant between the two sets of VCFs. Table 1 contains the counts of all genotypes in each concordance category, as well as definitions of each category. Among SNVs, primary QC of Atlas-generated SNV genotype data removed 2,605,141 low-quality variants (8.32%), while primary QC of GATK-generated SNV genotypes removed 3,493,548 low-quality variants (11.51%). For indels, primary QC among Atlas-generated genotypes removed 224,979 variants (5.85%) of low quality, and among GATK-generated indels removed 1,353,892 variants (27.82%). Once primary QC was completed, genotypes were compared between the two QCed VCFs.

Table 2 provides counts for the QC filtering of SNVs and indels at each step of variant-level QC (implemented after primary QC) as applied to Atlas and GATK VCFs. Overall, pipeline-specific QC filtering excluded an additional 1,932,851 SNVs (6.17%) from Atlas-generated SNV genotype data and 141,192 SNVs (0.46%) from GATK-generated SNV genotype data. QC filtering excluded 1,289,564 indels (30.91%) from Atlas and 362,593 indels (7.45%) from GATK. While most filtration steps excluded variants in both pipelines, it is notable that due to differences in processing of the BAMs by the centers, no variants with high read depth were identified among the GATK SNV or indel genotypes.

**Table 2.** WGS variant counts by QC filtration step. Entries in parentheses represent QC filters and the counts of variants excluded with the implementation of that filter, whereas non-parenthetical entries represent total counts after implementation of filtration steps.

| Counts | SNVs |  | Indels |  |
| --- | --- | --- | --- | --- |
|  | Atlas | GATK | Atlas | GATK |
| Total variants | 31,314,231 | 30,364,600 | 3,848,314 | 4,866,737 |
| (Multi-allelic variants) | (0) | (177,445) | (0) | (1,205,637) |
| Biallelic variants | 31,314,231 | 30,187,155 | 3,848,314 | 3,661,100 |
| Biallelic variants with<br>“PASS” (GATK) or<br>MS>0.8 (Atlas) | 28,709,090 | 26,871,052 | 3,623,335 | 3,512,845 |
| (Failed Monomorphic<br>filter) | (854,440) | (37,798) | (394,767) | (13,337) |
| (Failed Missing Rate filter) | (1,009,878) | (58,845) | (765,852) | (302,609) |
| (Failed Mean Depth filter) | (461) | (0) | (619) | (0) |
| (Failed Het filter<br>[MAF<0.2]) | (15,013) | (27,333) | (14,900) | (34,607) |
| (Failed Het filter<br>[MAF≥0.2]) | (53,059) | (17,216) | (13,336) | (12,040) |
| (Failed Het filter<br>[MAF≥0.2] due to all<br>heterozygous) | (31,131) | (0) | (0) | (0) |
| After variant QC | 26,776,239 | 26,729,860 | 2,433,771 | 3,150,252 |

For both SNVs and indels, a higher proportion of MIs was observed among GATK SNVs and indels (0.01087% and 0.08%, respectively) than among Atlas (0.00392% for SNVs and 0.02% for indels). This rate was highest among variants that were called only in the GATK VCF. For example, after pipeline-specific QC the MI rate for WGS SNVs was 0.12% for variants called only by GATK and 0.01% for variants called only by Atlas. A similar pattern among pipeline-unique variants was seen for indels (0.15% vs 0.04%). MIs in each VCF were counted but the genotypes implicated in the inconsistencies were retained. Table 3 provides a summary of MI rates from the pipeline specific QCed GATK and Atlas VCFs. Given the MI rates observed per parent-offspring pair (Figure 2), the sequence information appeared to be consistent with the relationships specified among the sequenced subjects in the pedigree files.

**Table 3.** Mendelian error rates from the QCed GATK and Atlas VCFs, and final post-QC consensus genotype set.

| Calling Pipeline | MI Category | SNVs | Indels |
| --- | --- | --- | --- |
| GATK | Total | 0.01087% | 0.08% |
|  | Unique | 0.12009% | 0.15% |
|  | Common | 0.00581% | 0.03% |
| Atlas | Total | 0.00392% | 0.02% |
|  | Unique | 0.01638% | 0.04% |
|  | Common | 0.00337% | 0.01% |
| Consensus | Total | 0.00420% | 0.02% |
|  | GATK unique | 0.02043% | 0.04% |
|  | Atlas unique | 0.01638% | 0.04% |
|  | Common | 0.01838% | 0.01% |

**Figure 2.**
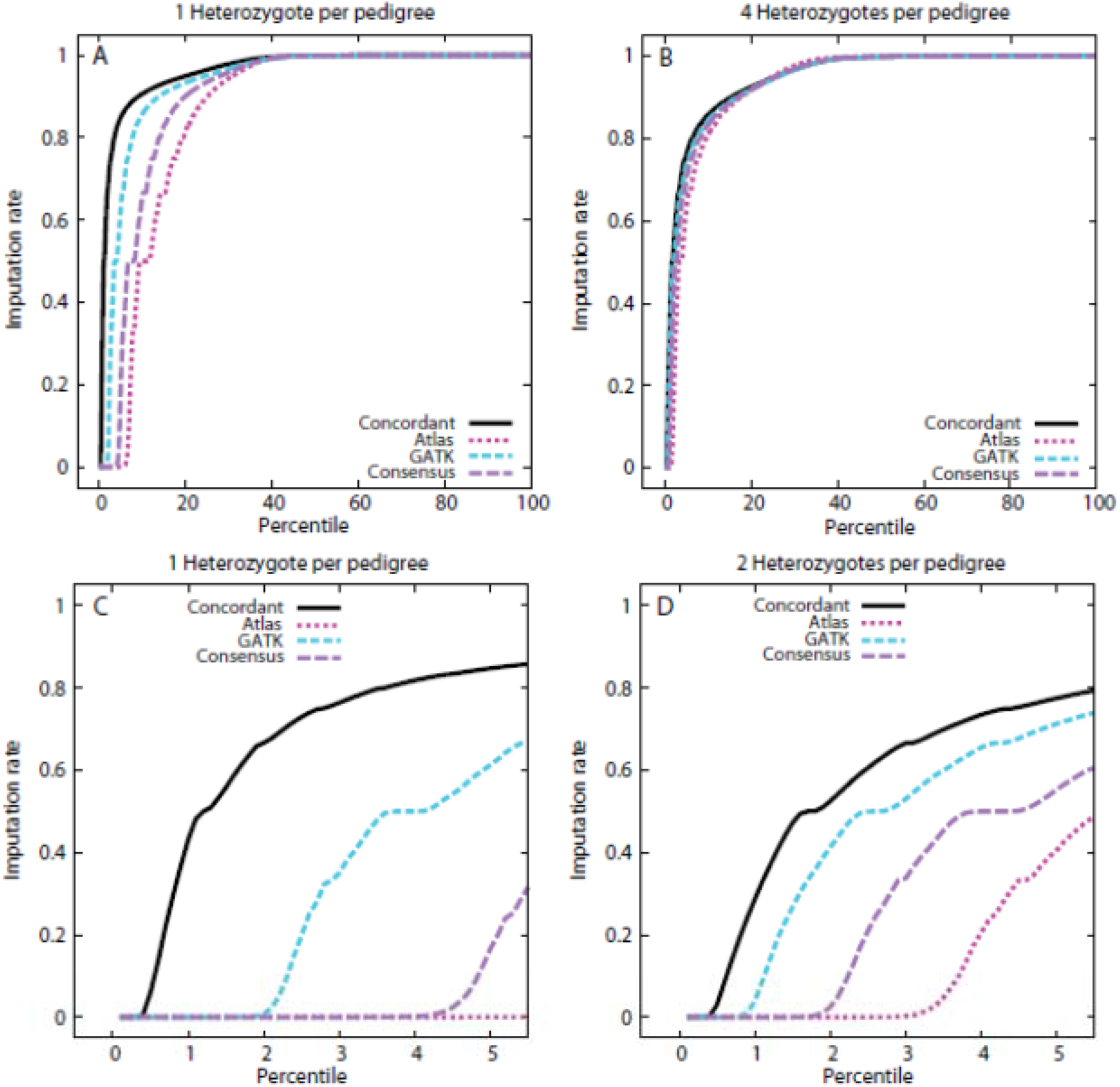
Cumulative distribution of genotype imputation rates for variants occurring 1-4 times as heterozygotes in the ADSP pedigrees. Panels A and B: full cumulative distributions for the cases of 1 (Panel A) and 4 (Panel B) heterozygotes per pedigree. Panels C and D provide detail for the variants with the lowest 5 percent of the imputation rate, for the cases of 1 (Panel C) and 2 (Panel D) heterozygotes per pedigree. The variants with observed data after the QC protocol to establish concordant calls between the pipelines are indicated by the solid black line (Concordant), while the variants subsequently retained as having high quality in one of the two pipelines are indicated by the long-dashed purple line (Consensus). The information for each of the two contributing pipelines to the Consensus variants is also represented, with the Atlas pipeline represented by the dark red, dotted line, and the GATK pipeline by the medium-dash blue line.

Table 4 provides the number of variants removed at each filtering step for the second round of post-consensus variant level QC. While pipeline-specific QC previously removed variants of low quality from the individual call sets, this second round of QC excluded variants that appeared low quality after consensus calling (e.g., after the exclusion of discordant genotypes, etc.). The frequency of concordant and pipeline-unique genotypes after consensus calling is summarized in Table 5. Table 6 provides the final number of variants and genotypes in the consensus WGS genotype sets.

**Table 4.** WGS variant counts with by post-consensus variant-level QC filtration step. Entries in parentheses represent counts of variants excluded with the implementation, whereas non-parenthetical entries represent total counts after implementation of filtration steps.

| Counts | SNVs | Indels |
| --- | --- | --- |
| Total variants | 27,970,909 | 3,549,344 |
| (Failed Monomorphic filter) | (12,058) | (166,069) |
| (Failed Missing Rate filter) | (61,252) | (244,045) |
| (Failed Excess Heterozygosity filter [MAF<0.2]) | (760) | (3,219) |
| (Failed Excess Heterozygosity filter [MAF≥0.2]) | (65) | (2,085) |
| Total after variant-level QC | 27,896,774 | 3,133,926 |

**Table 5.** Post pipeline-specific quality control (QC) percentages for genotype concordance, discordance, or calling in only one pipeline (Atlas or GATK).

| Genotype Pairs | SNVs | Indels |
| --- | --- | --- |
| Concordant | 90.93 | 55.62 |
| Discordant | 0.0107 | 0.286 |
| Called only in GATK | 4.56 | 27.99 |
| Called only in Atlas | 4.23 | 10.33 |
| Missing in both | 0.264 | 5.08 |
| Called in both with<br>different alternate allele | 0.000729 | 0.690 |

**Table 6.** Final counts of genotypes and variants after implementation of pipeline-specific QC, the consensus protocol, and post-consensus variant-level QC. “REF” indicates reference allele, and “ALT” indicates alternate allele.

| **Counts** | **SNVs** | **Indels** |
| --- | --- | --- |
| REF/REF | 14,050,706,093 | 1,595,662,048 |
| REF/ALT | 1,293,864,012 | 118,839,494 |
| ALT/ALT | 722,702,957 | 60,368,310 |
| Missing | 57,062,310 | 36,539,376 |
| **Total # variants** | **27,896,774** | **3,133,926** |

Mendelian inconsistencies were evaluated again in the final consensus genotype sets. Employing the consensus protocol reduced the MI rate among the variants called only by GATK from 0.12% to 0.02% in the SNVs and from 0.15% to 0.04% in the indels (see Table 3), providing evidence that the calls remaining in the consensus genotype set are likely higher quality.

### Data quality after implementation of QC and consensus calling

Most variants had extremely high imputation rates, with higher rates in the concordant than the consensus-only call sets (Figure 2, Table S4). The median imputation rate for variants was >0.995 for all configurations evaluated, and imputation rates at the lowest quartile were still high, ranging from 0.927 to 0.964 depending on the number of sequenced heterozygotes per pedigree. Even the very low fifth percentile achieved reasonably high imputation rates: for the concordant variants, imputation rates ranged from 0.762 to 0.848, while the consensus-only variant rates were 0.553-0.7 for families with 2-4 sequenced heterozygotes. A low imputation rate of 0.167 was found only for the consensus-only variants for families with a single heterozygote among the sequenced subjects. These observations suggest that the WGS genotypes are of high quality, overall, with only a small fraction that may have unusually high genotyping rates.

Genotypes in the concordant variant call set were of higher quality than those in the consensus call set. This inference is derived from the observation that imputation rates were higher, at all percentiles, for the concordant than consensus genotypes. The difference in quality between call sets was most extreme for variants heterozygous in only one sequenced subject (Figure 2A), with rapid attenuation of the difference with increasing numbers of heterozygotes per pedigree. Figure 2 panels A and B show results for the extremes of heterozygotes examined per pedigree. Intermediate results were obtained (not shown) for the remaining situations examined. The fraction of variants that had a 0% imputation rate (failed imputation) was also consistent with higher quality genotypes for the concordant variants. The fraction of concordant variants that failed imputation was a low 0.003-0.004 and was effectively independent of the number of heterozygotes per family. In contrast, for the consensus variants, the imputation failure rate was higher, and was a function of the number of heterozygotes per pedigree, decreasing monotonically from rates of 0.032 to 0.006 with increasing numbers of heterozygotes from one to four.

Genotypes in the consensus-only call set appeared to be of higher quality when derived from the GATK than the Atlas pipeline. The difference between the results for the two individual pipelines was most apparent in the case of single heterozygotes per pedigree, with the difference between pipelines particularly evident in the lowest tail of the distribution (Figure 2C-D). For variants with multiple heterozygotes per pedigree the differences between the two pipelines were minimal (Figure 2B). It is worth noting that of the variants that were in the consensus-only call set, the fraction of variants called by Atlas that could be found in the 1000 Genomes data^29^ (13.2%) was lower than the equivalent fraction for GATK (37.5%). Correspondingly there were lower allele frequencies for the Atlas variants than the GATK variants, with 55% and 50% of variants, respectively, below a 1000 Genomes EUR sample minor allele frequency of 0.05.

### Sample-level QC findings

Tables 7 and 8 present sample-level QC metrics, stratified by ethnicity and sequencing center, respectively. Few samples were flagged as outliers for sample-level QC metrics, suggesting high overall quality of all samples after QC. These samples were not excluded.

**Table 7.** Sample-level QC metrics stratified by ethnicity (EA, European American; ERF, Dutch Isolate; CH, Caribbean Hispanic). Values reported are mean, standard error (SE) in parentheses, and count of outliers at 4 SE and 6 SE, respectively, in square brackets unless otherwise specified.

| Sample-level Metrics | Ethnic Subgroup |  |  |
| --- | --- | --- | --- |
|  | EA | ERF | CH |
| # Samples (N) | 192 | 30 | 356 |
| # REF/REF genotypes | 24,484,692 (34,145) [1, 0] | 24,525,871 (20,686) [0, 0] | 24,190,270 (154,500) [0, 0] |
| # REF/ALT genotypes | 2,035,314 (23,228) [1, 0] | 2,023,070 (31,060) [0, 0] | 2,365,800 (161,320) [0, 0] |
| # ALT/ALT genotypes | 1,271,991 (10,318) [1, 0] | 1,284,034 (15,692) [0, 0] | 1,235,478 (32,757) [0, 0] |
| # Missing genotypes | 100,440 (43,356) [2, 0] | 59,462 (16,285) [0, 0] | 100,889 (42,421) [2, 2] |
| Missingness Rate (%) | 0.0036 [2, 0] | 0.0021 [0, 0] | 0.0036 [2, 2] |
| # Singletons | 7,005 (5,521) [0, 0] | 13,856 (5,030) [0, 0] | 8,158 (9,046) [2, 0] |
| # Doubletons | 8,467 (3,346) [0, 0] | 8,346 (1,854) [0, 0] | 12,567 (7,017) [0, 0] |
| Heterozygosity-to-Homozygosity ratio | 1.6003 (0.0256) [1, 0] | 1.576 (0.0421) [0, 0] | 1.9183 (0.1634) [0, 0] |
| Ti/Tv Ratio | 2.1256 (0.0027) [0,0] | 2.1242 (0.002) [0, 0] | 2.1239 (0.0032) [1, 0] |

**Table 8.** Sample-level QC metrics stratified by sequencing center. Values reported are mean, standard error (SE) in parentheses, and count of outliers at 4 SE and 6 SE, respectively, in square brackets unless otherwise specified.

| Sample-level Metrics | Sequencing Center |  |  |
| --- | --- | --- | --- |
|  | Broad | WashU | Baylor |
| # Samples (N) | 229 | 183 | 166 |
| # REF/REF genotypes [mean (SD)] | 24,275,244 (178,846) [0, 0] | 244,040,032 (176,288) [4, 0] | 24,238,582 (180,562) [0, 0] |
| # REF/ALT genotypes [mean (SD)] | 2,262,291 (196,200) [0, 0] | 2,127,330 (176,042) [0, 0] | 2,327,295 (196,027) [0, 0] |
| # ALT/ALT genotypes [mean (SD)] | 1,245,780 (31,703) [0, 0] | 1,266,015 (23,634) [0, 0] | 1,238,609 (35,480) [0, 0] |
| # Missing genotypes [mean (SD)] | 109,122 (42,151) [2, 0] | 95,060 (33,063) [3, 0] | 87,952 (49,452) [1, 1] |
| Missingness Rate (%) | 0.0039 [2, 0] | 0.0034 [3, 0] | 0.0032 [1, 1] |
| # Singletons [mean (SD)] | 7,662 (8,466) [1, 0] | 7,670 (7,155) [2, 0] | 9,076 (8,141) [0, 0] |
| # Doubletons [mean (SD)] | 10,976 (6,454) [0, 0] | 10,023 (5,598) [2, 0] | 12,062 (6,263) [0, 0] |
| Heterozygosity-to-Homozygosity ratio [mean (SD)] | 1.82 (0.1929) [0, 0] | 1.6824 (0.161) [0, 0] | 1.8842 (0.2029) [0, 0] |
| Ti/Tv Ratio [mean (SD)] | 2.1266 (0.0025) [0, 0] | 2.1238 (0.0024) [0, 0] | 2.1223 (0.0026) [1, 0] |

Figures S2 and S3 depict the distribution of heterozygosity for WGS samples after QC and consensus calling. Five members within one family were initially reported as European samples showed excess heterozygosity relative to other European samples, but were within the range of excess heterozygosity observed among Caribbean Hispanic samples (Figure S4). Later examination of clinical records found that the five samples were actually of African ancestry, and these were reclassified and co-analyzed with Caribbean Hispanic samples. The mean sample-specific genome-wide Ti/Tv ratio was 2.12 for all ethnic groups and sequencing centers (see Figures S2C and S3C and Tables 7 and 8). Whole genome sequencing is expected to have a Ti/Tv ratio of 2.10 for known variants^9^ and the consensus called variants fall at that expected threshold. Text S6 provides additional details on within-sample comparisons of three replicates done at each of the sequencing centers.

## Discussion

In the largest sequencing effort to date to discover rare genetic variation playing a direct role in AD, the ADSP has generated WGS data on 578 subjects from 111 multiplex AD families. Maximizing the potential to reveal true associations and minimizing potential false-positive findings requires comprehensive and rigorous quality control of sequence data prior to any analysis. In order to provide a consistent high-quality dataset, the QC working group of the ADSP developed and implemented a novel QC protocol that generated a concordant genotype set and a consensus genotype set from two variant calling pipelines. These QCed datasets are available via the database of Genotypes and Phenotypes (dbGaP) and the NIA Genetics of Alzheimer’s Disease Data Storage Site (NIAGADS) portal and include PLINK binary files and VCF files containing only genotypes that passed QC and QC annotation files containing information on all variants regardless of pass/fail status. Use of these standardized QCed datasets will also facilitate comparison across analyses by different research groups, efficiencies that are important to aid in the discovery of AD-related genetic factors. It is anticipated that this information will streamline future *in silico* replication using ADSP data. The availability of these datasets will increase efficiency in the use of the ADSP data by precluding the need for each authorized investigator to perform QC.

Among the most useful features for data users in this QC process is the creation of ‘companion’ files, or QC annotation files. These files provide a dataset-specific reference guide containing information on all variants and genotypes called by either pipeline regardless of whether the variants passed or failed and include the pass/fail status for all original called variants after QC filters are applied. This detailed information (Table S3) on the disposition of all variants serves multiple purposes. Firstly, it provides a detailed record of all quality issues identified during the QC process, improving reproducibility of the process. Secondly, it allows for comprehensive examinations of widely agreed-upon QC filters and their downstream effects on data quality; by utilizing annotation that identifies all QC criteria for which a variant fails independently of order of implementation, authorized investigators can know the effects of applying any QC filters in any desired order before applying them. Thirdly, the QC annotation can be used to confirm that the QC process has been implemented correctly, as recorded values of metrics on failing variants should be consistent with exclusion criteria. Finally, these files explain why variants of interest may be missing from the QCed data. Typically, information on variants that are removed is not recorded, and which QC criteria a variant may fail are subject to the order of QC filter implementation. Notably, our approach identifies all QC criteria a variant may fail, which makes it independent of QC filter implementation order and fully reproducible.

One of the strengths of the study, yet one that provided the biggest challenge for QC, was the use of multiple variant-calling pipelines (Atlas and GATK). Different variant-calling algorithms have been developed and the strengths and weaknesses of each are not well documented. Elucidating specific conditions (e.g., what type of variant or sequencing method) under which each algorithm performs optimally is beyond the scope of the current study. The current goal was to use the information from the two calling pipelines to generate a set of consistently high-quality genotypes to facilitate the discovery of AD-related genetic variants.

The concordant data set includes genotypes from variants that were called by both pipelines, met all filtering criteria, and were identical between the two pipelines. For SNVs and indels, 91% and 56% of genotypes, respectively, were concordant between the two pipelines. These concordant calls represent our highest-confident genotype set. The quality of these concordant calls was further supported by the lower MI rate and higher mean GQ levels relative to discordant calls.

The concordant genotype set does not, however, take advantage of unique strengths of the two calling algorithms applied and may be overly conservative. To address this issue, a consensus protocol was developed that incorporates variants called by only one of the pipelines. For WGS SNVs and indels, a larger number of variants uniquely called by GATK compared to Atlas were included in the consensus sets. Examination of MI indicated that after implementation of the pipeline-specific QC filters that the GATK-only variants might be more error prone than variants called only in Atlas or by both pipelines. This is likely driven by the fact that the pipeline-specific QC protocol for Atlas included both genotype-specific and variant-specific filtering, whereas the pipeline specific QC protocol for GATK included only variant-specific filtering, following sequencing center recommendations. Implementation of a GQ filter on genotypes within variants called only by GATK improved the quality as measured by MI rate. The final consensus genotype datasets include the genotypes that were concordant in both pipelines, and additionally include high-quality genotypes from variants called in only one of the two pipelines. While concordant variants are of generally higher quality than the high-quality genotypes called in only one pipeline, it should be noted that calling criteria or artifacts in one pipeline may lead to true causal variants being missed in that pipeline, and this approach maximizes the number of high-quality variants retained to improve the likelihood of identifying causal variants in a dataset.

The Mendelian consistency rate in family-based imputation was used to compare the quality of genotype calls when two calling algorithms were implemented. A first question addressed was whether use of two pipelines resulted in improved quality of variants called relative to use of only a single pipeline; a second was whether there was a difference in the quality of variants called between the two individual pipelines. Although not specifically designed for the purpose used here, the MC rate during pedigree-based imputation makes use of results of other ongoing computations (e.g., generation of inheritance vectors in pedigrees and MI frequency) and was sufficient to address the two questions of interest. These results suggest that the genotypes called identically in both pipelines were of higher quality than calls from a single pipeline. While the consistency rate was slightly higher for variants called only by GATK compared to those called only by Atlas, this difference was very small and overall quality after pipeline-specific QC was high for both Atlas- and GATK-generated genotypes.

The QC protocol implemented on the ADSP WGS data is generalizable to other large-scale sequencing studies. While the protocol implemented for this study utilized data-driven thresholds and hence specific values (e.g., GQ thresholds or heterozygosity score statistic threshold) that may not translate to other studies, the pipeline protocol simplifies determination of these threshold for each novel dataset, tailoring the pipeline to the unique characteristics of the dataset and allowing for easy implementation. This protocol offers a paradigm for other studies to develop their own metrics and, under the condition of using multiple genotype calling pipelines, an approach not previously applied to large NGS datasets. As more sequencing studies are conducted, the potential for universal guidelines may be feasible, and many features of the ADSP pipeline, such as QC filtering on multiple criteria in parallel, have been designed with adaptability to future guidelines in mind. For example, the pipeline was constructed to allow for the assessment of HWE using tests that assume independence of observations are part of the variant-level QC if most or all samples in the dataset are unrelated or using empirical thresholds with an excess heterozygosity statistic as was done here with family data. It should be noted that in population-based datasets, assessments of quality would be unable to utilize MI checking unless some informative relative pairs are included in sequencing to examine this quality; alternative quality assessments could be applied to similar effect, including genotype concordance between sequence and GWAS data by strata of GQ values, concordance among unfiltered variants across replicate samples, etc.

This work also highlights the value of utilizing information available prior to sequencing, including identification of parent-offspring pairs and high-density genotyping chip data. Even though identification of MIs is limited with biallelic variants, the change in MI rate with different genotype sets provided important information on genotype quality. Inclusion of relative pairs for QC may be useful even if the main study design is based on unrelated cases and controls. Similarly, the availability of GWAS data for sample validation was also important, both in determining quality of sequence genotypes through concordance checks, and in characterizing between-sample relatedness and population substructure with a high quality subset of GWAS genotypes. For these reasons, the availability of additional data from sequenced samples, such as relatedness information and the availability of other independent genetic resources, should be considered when prioritizing samples for sequencing and/or designing sequencing projects.

In summary, the ADSP has generated high-quality QCed WGS datasets by developing and implementing a novel QC protocol that integrates calls from both the Atlas and GATK variant calling algorithms. This approach provides a model for QC for other large-scale sequencing studies. The distribution of these carefully QCed high-quality WGS SNV and indel genotype sets, shared via the Database for Genotypes and Phenotypes (dbGaP), will provide an important public resource for untangling the genetic etiology of AD.

## Supplemental Data

**Table S1.** Sample counts by sequencing center

**Text S1.** Library preparation and whole genome sequencing

**Table S2.** Parameters used for whole genome alignment and calling by sequencing center

**Text S2.** Description and workflows for genotyping calling in GATK and Atlas

**Text S3.** Filtering for Extreme Heterozygosity

**Text S4.** Determination of GQ thresholds for GATK genotype filtering

**Table S3.** QC Annotation file field descriptions for variant-level QC and sample-level QC

**Text S5.** Estimation of Inconsistency Rate in Family-based Imputed Genotypes

**Table S4.** Mean imputation rates across pedigrees

**Text S6.** Comparison of samples replicated between sequencing centers

**Table S5.** Concordance among replicate sample genotypes from GATK

**Table S6.** Concordance among replicate sample genotypes from Atlas

**Figure S1.** Potential genotype configurations where there are Mendelian inconsistencies (MI) and discordant calls between the two pipelines.

**Figure S2.** Distribution of the heterozygosity-to-homozygosity ratio among WGS samples prior to variant-level QC and consensus calling, stratified by race/ethnicity.

**Figure S3.** Distributions of sample-level metrics among WGS samples after variant-level QC and consensus calling, stratified by race/ethnicity

**Figure S4.** Distributions of sample-level metrics among WGS samples after variant-level QC and consensus calling, stratified by sequencing center

## References

1. Pareek, C.S., Smoczynski, R. & Tretyn, A. Sequencing technologies and genome sequencing. J Appl Genet 52, 413–35 (2011). (PMID: 21698376/Standin: 3189340)

2. Zhou, Q., Su, X., Wang, A., Xu, J. & Ning, K. QC-Chain: fast and holistic quality control method for next-generation sequencing data. PLoS One 8, e60234 (2013). (PMID: 23565205/Standin: 3615005)

3. Guo, Y., Ye, F., Sheng, Q., Clark, T. & Samuels, D.C. Three-stage quality control strategies for DNA re-sequencing data. Brief Bioinform 15, 879–89 (2014). (PMID: 24067931/Standin: 4492405)

4. Patel, R.K. & Jain, M. NGS QC Toolkit: a toolkit for quality control of next generation sequencing data. PLoS One 7, e30619 (2012). (PMID: 22312429/Standin: 3270013)

5. Schmieder, R. & Edwards, R. Quality control and preprocessing of metagenomic datasets. Bioinformatics 27, 863–4 (2011). (PMID: 21278185/Standin: 3051327)

6. Li, B., Zhan, X., Wing, M.K., Anderson, P., Kang, H.M. & Abecasis, G.R. QPLOT: a quality assessment tool for next generation sequencing data. Biomed Res Int 2013, 865181 (2013). (PMID: 24319692/Standin: 3844194)

7. Guo, Y., Zhao, S., Sheng, Q., Ye, F., Li, J., Lehmann, B., Pietenpol, J., Samuels, D.C. & Shyr, Y. Multi-perspective quality control of Illumina exome sequencing data using QC3. Genomics 103, 323–8 (2014). (PMID: 24703969/Standin:

8. McKenna, A., Hanna, M., Banks, E., Sivachenko, A., Cibulskis, K., Kernytsky, A., Garimella, K., Altshuler, D., Gabriel, S., Daly, M. & DePristo, M.A. The Genome Analysis Toolkit: a MapReduce framework for analyzing next-generation DNA sequencing data. Genome Res 20, 1297–303 (2010). (PMID: 20644199/Standin: 2928508)

9. DePristo, M.A., Banks, E., Poplin, R., Garimella, K.V., Maguire, J.R., Hartl, C., Philippakis, A.A., del Angel, G., Rivas, M.A., Hanna, M., McKenna, A., Fennell, T.J., Kernytsky, A.M., Sivachenko, A.Y., Cibulskis, K., Gabriel, S.B., Altshuler, D. & Daly, M.J. A framework for variation discovery and genotyping using next-generation DNA sequencing data. Nat Genet 43, 491–8 (2011). (PMID: 21478889/Standin: 3083463)

10. Van der Auwera, G.A., Carneiro, M.O., Hartl, C., Poplin, R., Del Angel, G., Levy-Moonshine, A., Jordan, T., Shakir, K., Roazen, D., Thibault, J., Banks, E., Garimella, K.V., Altshuler, D., Gabriel, S. & DePristo, M.A. From FastQ data to high confidence variant calls: the Genome Analysis Toolkit best practices pipeline. Curr Protoc Bioinformatics 43, 11 10 1–33 (2013). (PMID: 25431634/Standin: 4243306)

11. Challis, D., Yu, J., Evani, U.S., Jackson, A.R., Paithankar, S., Coarfa, C., Milosavljevic, A., Gibbs, R.A. & Yu, F. An integrative variant analysis suite for whole exome next-generation sequencing data. BMC Bioinformatics 13, 8 (2012). (PMID: 22239737/Standin: 3292476)

12. Morrison, A.C., Voorman, A., Johnson, A.D., Liu, X., Yu, J., Li, A., Muzny, D., Yu, F., Rice, K., Zhu, C., Bis, J., Heiss, G., O'Donnell, C.J., Psaty, B.M., Cupples, L.A., Gibbs, R., Boerwinkle, E., Cohorts for, H. & Aging Research in Genetic Epidemiology, C. Whole-genome sequence-based analysis of high-density lipoprotein cholesterol. Nat Genet 45, 899–901 (2013). (PMID: 23770607/Standin: 4030301)

13. Kunkle, B.W., Jaworski, J., Barral, S., Vardarajan, B., Beecham, G.W., Martin, E.R., Cantwell, L.S., Partch, A., Bird, T.D., Raskind, W.H., DeStefano, A.L., Carney, R.M., Cuccaro, M., Vance, J.M., Farrer, L.A., Goate, A.M., Foroud, T., Mayeux, R.P., Schellenberg, G.D., Haines, J.L. & Pericak-Vance, M.A. Genome-wide linkage analyses of non-Hispanic white families identify novel loci for familial late-onset Alzheimer’s disease. Alzheimers Dement 12, 2–10 (2016). (PMID: 26365416/Standin: 4717829)

14. Barral, S., Cheng, R., Reitz, C., Vardarajan, B., Lee, J., Kunkle, B., Beecham, G., Cantwell, L.S., Pericak-Vance, M.A., Farrer, L.A., Haines, J.L., Goate, A.M., Foroud, T., Boerwinkle, E., Schellenberg, G.D. & Mayeux, R. Linkage analyses in Caribbean Hispanic families identify novel loci associated with familial late-onset Alzheimer’s disease. Alzheimers Dement 11, 1397–406 (2015). (PMID: 26433351/Standin: 4690771)

15. Beecham, G.W., Bis, J.C., Martin, E.R., Choi, S.-H., DeStefano, A.L., van Duijn, C.M., Fornage, M., Gabriel, S.B., Koboldt, D.C., Larson, D.E., Naj, A.C., Psaty, B.M., Salerno, W., Bush, W.S., Foroud, T.M., Wijsman, E., Farrer, L.A., Goate, A., Haines, J.L., Pericak-Vance, M.A., Boerwinkle, E., Mayeux, R., Seshadri, S. & Schellenberg, G.D. The Alzheimer’s Disease Sequencing Project: study design and sample selection. (2017). (PMID:

16. Liu, F., Arias-Vasquez, A., Sleegers, K., Aulchenko, Y.S., Kayser, M., Sanchez-Juan, P., Feng, B.J., Bertoli-Avella, A.M., van Swieten, J., Axenovich, T.I., Heutink, P., van Broeckhoven, C., Oostra, B.A. & van Duijn, C.M. A genomewide screen for late-onset Alzheimer disease in a genetically isolated Dutch population. Am J Hum Genet 81, 17–31 (2007). (PMID: 17564960/Standin: 1950931)

17. Nato, A.Q., Jr., Chapman, N.H., Sohi, H.K., Nguyen, H.D., Brkanac, Z. & Wijsman, E.M. PBAP: a pipeline for file processing and quality control of pedigree data with dense genetic markers. Bioinformatics 31, 3790–8 (2015). (PMID: 26231429/Standin: 4668752)

18. O'Connell, J.R. & Weeks, D.E. PedCheck: a program for identification of genotype incompatibilities in linkage analysis. Am J Hum Genet 63, 259–66 (1998). (PMID: 9634505/Standin: 1377228)

19. Trubetskoy, V., Rodriguez, A., Dave, U., Campbell, N., Crawford, E.L., Cook, E.H., Sutcliffe, J.S., Foster, I., Madduri, R., Cox, N.J. & Davis, L.K. Consensus Genotyper for Exome Sequencing (CGES): improving the quality of exome variant genotypes. Bioinformatics 31, 187–93 (2015). (PMID: 25270638/Standin: 4287941)

20. Zook, J.M., Chapman, B., Wang, J., Mittelman, D., Hofmann, O., Hide, W. & Salit, M. Integrating human sequence data sets provides a resource of benchmark SNP and indel genotype calls. Nat Biotechnol 32, 246–51 (2014). (PMID: 24531798/Standin:

21. Cantarel, B.L., Weaver, D., McNeill, N., Zhang, J., Mackey, A.J. & Reese, J. BAYSIC: a Bayesian method for combining sets of genome variants with improved specificity and sensitivity. BMC Bioinformatics 15, 104 (2014). (PMID: 24725768/Standin: 3999887)

22. Patel, Z.H., Kottyan, L.C., Lazaro, S., Williams, M.S., Ledbetter, D.H., Tromp, H., Rupert, A., Kohram, M., Wagner, M., Husami, A., Qian, Y., Valencia, C.A., Zhang, K., Hostetter, M.K., Harley, J.B. & Kaufman, K.M. The struggle to find reliable results in exome sequencing data: filtering out Mendelian errors. Front Genet 5, 16 (2014). (PMID: 24575121/Standin: 3921572)

23. Wall, J.D., Tang, L.F., Zerbe, B., Kvale, M.N., Kwok, P.Y., Schaefer, C. & Risch, N. Estimating genotype error rates from high-coverage next-generation sequence data. Genome Res 24, 1734–9 (2014). (PMID: 25304867/Standin: 4216915)

24. Carson, A.R., Smith, E.N., Matsui, H., Braekkan, S.K., Jepsen, K., Hansen, J.B. & Frazer, K.A. Effective filtering strategies to improve data quality from population-based whole exome sequencing studies. BMC Bioinformatics 15, 125 (2014). (PMID: 24884706/Standin: 4098776)

25. De Summa, S., Malerba, G., Pinto, R., Mori, A., Mijatovic, V. & Tommasi, S. GATK hard filtering: tunable parameters to improve variant calling for next generation sequencing targeted gene panel data. BMC Bioinformatics 18, 119 (2017). (PMID: 28361668/Standin: 5374681)

26. Ewels, P., Magnusson, M., Lundin, S. & Kaller, M. MultiQC: summarize analysis results for multiple tools and samples in a single report. Bioinformatics 32, 3047–8 (2016). (PMID: 27312411/Standin: 5039924)

27. Lander, E.S. & Green, P. Construction of multilocus genetic linkage maps in humans. Proc Natl Acad Sci U S A 84, 2363–7 (1987). (PMID: 3470801/Standin: 304651)

28. Cheung, C.Y., Thompson, E.A. & Wijsman, E.M. GIGI: an approach to effective imputation of dense genotypes on large pedigrees. Am J Hum Genet 92, 504–16 (2013). (PMID: 23561844/Standin: 3617386)

29. Genomes Project, C., Abecasis, G.R., Altshuler, D., Auton, A., Brooks, L.D., Durbin, R.M., Gibbs, R.A., Hurles, M.E. & McVean, G.A. A map of human genome variation from population-scale sequencing. Nature 467, 1061–73 (2010). (PMID: 20981092/Standin: 3042601)

